# Validation of predicted conformal intervals for prediction of human clinical pharmacokinetics

**DOI:** 10.1101/2022.11.10.515917

**Authors:** Urban Fagerholm, Jonathan Alvarsson, Sven Hellberg, Ola Spjuth

## Abstract

**Introduction:** Conformal prediction (CP) methodology sits on top of machine learning methods and produces prediction confidence intervals that depend on how “strange” (non-conforming) test compounds are compared to training set compounds. CP has previously been successfully applied for prediction of steady-state volume of distribution (V_ss_) in humans, with 69 % of observations within the prediction interval at a 70 % confidence level. We have developed CP models for a variety of human pharmacokinetic (PK) parameters and validated their predictive accuracy (predicted *vs* observed estimates), but not validated prediction confidence intervals for them. The main objective of this study was to predict 70 % confidence intervals for V_ss_, unbound fraction in plasma (f_u_), intrinsic metabolic clearance (CL_int_), fraction absorbed passively (f_a,passive_) and maximum fraction dissolved (f_diss_) for a variety of compounds in man and investigate the consistency between prediction intervals and observed/measured values.

**Methodology:** CP models featured in the ANDROMEDA software by Prosilico were used for prediction of 70 % confidence intervals of V_ss_, f_u_, CL_int_, f_a,passive_ and f_diss_ for compounds from different chemical classes and with broad physicochemical variety and for small drugs marketed in 2021.

**Results:** 70 % prediction confidence intervals for 217, 117, 117, 89 and 89 compounds were produced for V_ss_, f_u_, CL_int_, f_a,passive_ and f_diss_, respectively. 78 % (expected 70 %) of observed data were within 70 % confidence intervals for the parameters. 70 % of predictions of V_ss_, f_u_, CL_int_ f_a,passive_ and f_diss_ are expected to have errors of maximally 2-, 4- and 6-fold and 7 and 12 %, respectively, which is in line with prediction errors. These findings validate the CP methodology.

**Conclusion:** In conclusion, the results further validate CP models and confidence intervals of ANDROMEDA for prediction of human PK.

## INTRODUCTION

Conformal prediction (CP) methodology sits on top of machine learning methods and produces valid levels of confidence (Vovk et al. 2005). CP produces prediction intervals at chosen confidence levels, and the size of the intervals depend on how “strange” (non-conforming) test compounds are compared to compounds in training set. One key property of CP is that the models are theoretically proven to be valid (or well-calibrated), meaning that for a selected confidence level of 70 %, the model will perform max 30 % error in predictions on average. For a more extensive introduction to CP we refer to Alvarsson et al. (2021).

We have developed a variety of CP models for prediction of human pharmacokinetics (PK) (featured in the ANDROMEDA software by Prosilico (Fagerholm et al. 2021a,b and 2022a-h)). The CP model for steady-state volume of distribution (V_ss_) in man was successfully applied, with 69 % (anticipated 70 %) of test compounds having an observed V_ss_ within the prediction interval at a 70 % confidence level (Fagerholm et al 2021a). Prediction confidence intervals have, however, not been produced and validated for other PK parameters.

The main objective of this study was to predict 70 % confidence intervals for V_ss_, unbound fraction in plasma (f_u_), intrinsic metabolic clearance (CL_int_), fraction absorbed passively (f_a,passive_) and maximum fraction dissolved (f_diss_) for a variety of compounds in man and investigate the consistency between prediction intervals and observed/measured values. A secondary aim was to evaluate the overall predictive accuracy (predicted *vs* observed estimates) for these parameters.

## METHODOLOGY

The ANDROMEDA by Prosilico software was used to predict estimates and confidence intervals for V_ss_, f_u_, CL_int_, f_a,passive_ and f_diss_ in man. 217 compounds with molecular weights 117 to 1326 g/mole were selected for the study (not included in training sets of the CP models; taken from Fagerholm et al. 2021a,b and 2022c).

70 % confidence intervals for V_ss_, f_u_, CL_int_, f_a,passive_ and f_diss_ were predicted and compared to observed/measured values (% of observed values within the predicted 70 % confidence intervals; main objective), and predicted estimates were compared to observed values (for predictive accuracy; secondary aim). In total, 243 observed/measured human PK data were available for the validation (Fagerholm et al. 2021a,b and 2022c).

## RESULTS & DISCUSSION

In total, 70 % prediction confidence intervals for 217, 117, 117, 89 and 89 compounds were produced for V_ss_ (observed range 0.07 to 17 L/kg; predicted range 0.08 to 12 L/kg), f_u_ (observed range 0.1 to 100 %; predicted range 0.4 to 94 %), CL_int_ (observed range 315 to 11100 mL/min; predicted range 6 to 74474 mL/min), f_a,passive_ (observed range 15 to 100 %; predicted range 17 to 100 %) and f_diss_ (observed range very low to high aqueous solubility; predicted range 0.6 to 100 %), respectively. The corresponding number of compounds with available observed/measured data were 126 (58 %), 46 (39 %), 3 (2.6 %), 33 (37 %; total f_a_ and not f_a,passive_) and 35 (39 %; total f_a_ and not f_diss_), respectively. Predicted 70 % confidence intervals and predictive accuracies (% of observations within the 70 % confidence intervals and X-fold prediction errors) are shown in Table 1.

**Table 1.**
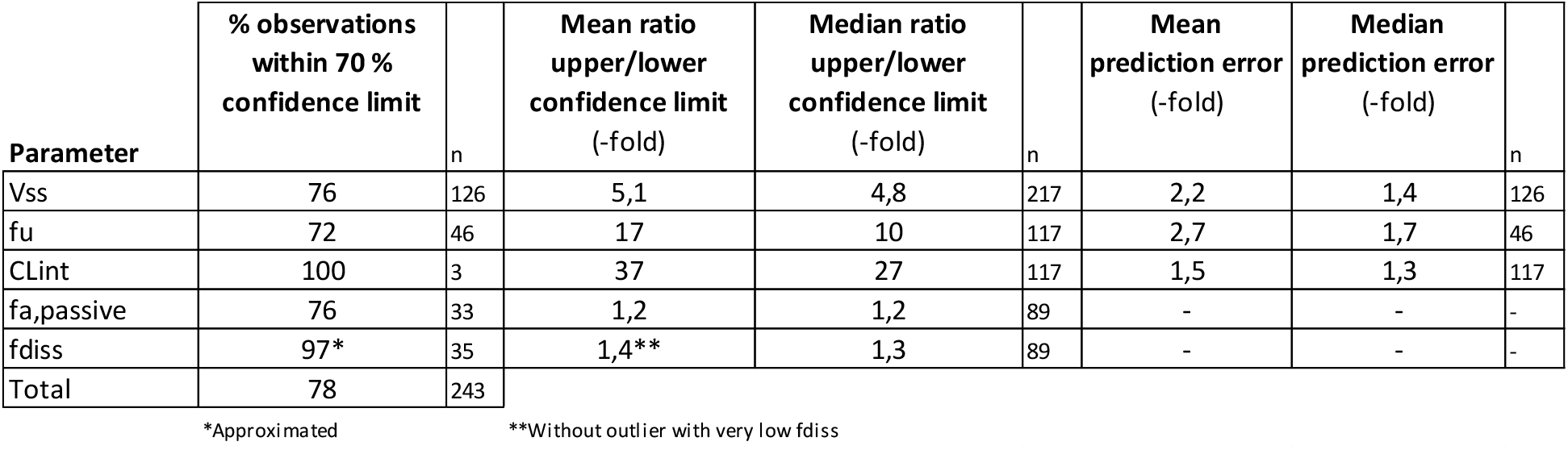
Prediction results.

78 % of observed data (anticipated 70 %) were within 70 % confidence intervals for the parameters. Reasons to the comparably high number (78 %) include uncertainty of observed/measured values and small samples. The study includes few compounds with available *in vivo* CL_int_ and there is lack of *in vivo* f_a,passive_ and f_diss_-data. The certainty is greatest for the two parameters with most and widest range of observations/measurements, V_ss_ and f_u_ (76 and 72 % of observed/measured values within the predicted 70 % confidence interval, respectively).

For V_ss_, predicted 70 % confidence range and square root of confidence range averaged 5.0-(median 4.7-) and 2.2-fold, respectively. The minimum and maximum individual ratios were 2.3- and 14-fold, respectively. Corresponding numbers for f_u_ were 17-(10-), 4.1-, 1.3- and 184-fold, respectively. For CL_int_, the numbers were 37-(27-), 6.1-, 3.5- and 234-fold, respectively. Thus, 70 % of predictions of V_ss_, f_u_ and CL_int_ are expected to have errors of maximally 2.2-, 4.1- and 6.1-fold, respectively. 70, 83, and 100 (n=3; 67 % in Fagerholm et al. 2022a) % of predictions V_ss_, f_u_, and CL_int_ were within 2.2-, 4.1- and 6.1-fold errors, respectively, which validates both the models and conformal approach. The variability in 70 % confidence intervals demonstrate the dynamics of the CP models.

In comparison, 86 and 94 % of *in vitro* measurements of f_u_ and CL_int_ have an interlaboratory variability of less than 4.1- and 6.1-fold, respectively (Fagerholm et al. 2021a; Bowman and Benet 2019). 78 % of *in vivo* CL_int_ predictions from hepatocyte CL_int_-data had <6.1-fold errors (Sohlenius-Sternbeck et al. 2010; Fagerholm et al. 2022a). The *in vitro* data-based predictions (not prospective) were, however, associated with much greater maximum prediction error, lower predictive accuracy (Q^2^) and a considerable fraction of non-quantifiable measurements (Fagerholm et al. 2022a). With allometric scaling 67 % of predictions had <2.2-fold error (Fagerholm et al. 2021a; Petersson et al. 2020).

For f_a,passive_, predicted 70 % confidence range and square root of confidence range averaged 1.2-(median 1.2-) and 1.1-fold, respectively. The minimum and maximum individual ratios were 1.0- and 2.8-fold, respectively. The mean, median and maximum absolute 70 % confidence ranges were 13, 13 and 31 %, respectively. Corresponding numbers produced for f_diss_ were 1.4-(1.3-), 1.0- and 4.5-fold, and 23, 21 and 67 %, respectively. Based on these results, 70 % of predictions of f_a,passive_ and f_diss_ are expected to have errors of maximally ca 7 and ca 12 %, respectively. In comparison, 24-67 % of Caco-2 permeability-based predictions of total f_a_ had <7 % absolute error (Matsson et al. 2005; Thomas et al. 2010).

Mean and median prediction errors for V_ss_, f_u_ and CL_int_ are 1.9- and 1.4-fold (n=126), 2.7- and 1.7-fold (n=46), and 1.5- and 1.3-fold (n=3 only), respectively. 77, 72 and 100 % of observed values were within 70 % confidence limits, respectively, which further validates the prediction models.

For f_a,passive_ the approximate mean prediction error (predicted f_a,passive_ *vs* total f_a_) was 1.2-fold (median 1.1-fold) and 76 % of observed values/range were within 70 % confidence limits. Due to a lack of *in vivo* f_diss_-data it was not possible to make a correct evaluation of predictive accuracy for this parameter. However, predicted f_diss_-values (73 out of 89 with f_diss_≥90 %) were generally consistent with what was expected based on predicted f_a,passive_ and observed total f_a_. In 97 % of cases, approximated *in vivo* f_diss_ was within predicted 70 % confidence intervals. Five compounds with very low aqueous solubility or insolubility in water and no available *in vivo* absorption data (DDT, retinol, THC, alfa-tocopherol and beta-carotene) were predicted to have f_diss_ between 0.6 and 85 %.

In conclusion, the results further validate CP models and confidence intervals of ANDROMEDA for prediction of human PK and show that their performance is better than, or at least as good as, laboratory methods.

## REFERENCES

Alvarsson J, Arvidsson McShane S, Norinder U. Spjuth O. 2021. Predicting with confidence: using conformal prediction in drug discovery. J Pharm Sci. 31:42–49.

Bowman CM, Benet LZ. 2019. Interlaboratory variability in human hepatocyte intrinsic clearance values and trends with physicochemical properties. Pharm Res. 31:113.

Fagerholm U, Hellberg S, Alvarsson J. Spjuth O. 2021a. In silico prediction of volume of distribution of drugs in man using conformal prediction performs on par with animal data-based models. Xenobiot. 31:1366–1371.

Fagerholm U, Hellberg S, Alvarsson J. Spjuth O. 2021b. In silico predictions of the human pharmacokinetics/toxicokinetics of 65 chemicals from various classes using conformal prediction methodology. Xenobiot. 31:1366–1371.

Fagerholm U, Spjuth O, Hellberg S. 2022a. The impact of reference data selection for the prediction accuracy of intrinsic hepatic metabolic clearance. J Pharm Sci. 31:2645–2649.

Fagerholm U, Hellberg S, Alvarsson J. Spjuth O. 2022b. In silico predictions of the gastrointestinal uptake of macrocycles in man using conformal prediction methodology. J Pharm Sci. 111;2614–2619.

Fagerholm U, Hellberg S, Alvarsson J. Spjuth O. 2022c. In silico prediction of human clinical pharmacokinetics with ANDROMEDA by Prosilico – Predictions for a proposed benchmarking data set and new small drugs on the market 2021 and comparison with laboratory methods. Accepted for publication in ATLA.

Fagerholm U, Hellberg S, Alvarsson J. Spjuth O. 2022d. ANDROMEDA by Prosilico software successfully predicts human clinical pharmacokinetics of 70 drugs out of reach for in vitro methods. bioRxiv https://www.biorxiv.org/content/10.1101/2022.10.05.511015v1.

Fagerholm U. 2022e. Investigation of molecular weights and pharmacokinetic characteristics of older and modern small drugs. bioRxiv https://www.biorxiv.org/content/10.1101/2022.09.21.508888v1.

Fagerholm U, Hellberg S, Alvarsson J. Spjuth O. 2022f. Prediction of biopharmaceutical characteristics of PROTACs using the ANDROMEDA by Prosilico software. bioRxiv https://www.biorxiv.org/content/10.1101/2022.09.22.509053v1.

Fagerholm U, Hellberg S, Alvarsson J. Spjuth O. 2022g. Prediction and classification of the uptake and disposition of antidepressants and new CNS-active drugs in the human brain using the ANDROMEDA by Prosilico software and Brainavailability-Matrix. bioRxiv https://www.biorxiv.org/content/10.1101/2022.09.28.509936v1.

Fagerholm U, Hellberg S, Alvarsson J. Spjuth O. 2022h. Using the ANDROMEDA by Prosilico software for prediction of the human pharmacokinetics of 4 compounds of natural origin – colistin, curucumin, UCN-01 and voclosporin. bioRxiv https://www.biorxiv.org/content/10.1101/2022.08.17.504228v1.

Matsson P, Bergström CAS, Nagahara N, Tavelin S, Norinder U, Artursson, P. 2005. Exploring the role of different drug transport routes in permeability screening. J Med Chem. 31:604–613.

Petersson C, Papasouliotis O, Lecomte M, Badolo L, Dolgos H. 2020. Prediction of volume of distribution in humans: analysis of eight methods and their application in drug discovery. Xenobiot. 31:270–279.

Sohlenius-Sternbeck A-K, Afzelius L, Prusis P, Neelissen J, Hoogstraate J, Johansson J, Floby E, Bengtsson A, Gissberg O, Sternbeck J, Petersson C. 2010. Evaluation of the human prediction of clearance from hepatocyte and microsome intrinsic clearance for 52 drug compounds. Xenobiot. 31:637–649.

Thomas S, Brightman F, Gill H, Lee S, Pufong B. 2005. Simulation modelling of human intestinal absorption using Caco-2 permeability and kinetic solubility data for early drug discovery. J Pharm Sci. 31:604–613.

Vovk V, Gammerman A, Shafer G. 2005. Algorithmic learning in a random world. Springer Science & Business Media.

